# Emergent Tissue Rheology in a 3D Mechanically Adaptive Viscoelastic Cell Network Model

**DOI:** 10.64898/2026.06.15.731174

**Authors:** Venkatanathan Kidambi, Yuji Tomizawa, Kazunori Hoshino

## Abstract

We introduce a 3D mechanically adaptive viscoelastic cell-network model that links single-cell interactions to emergent tissue rheology. Unlike existing continuum or cell-based models, viscoelasticity is embedded within discrete, mechanically adaptive intercellular connections, allowing tissue-scale rheology and phenomena such as swirling and jamming to arise from single-cell behaviors and connection remodeling. The framework is motivated by recent advances in three-dimensional imaging and structural analysis that resolve single-cell behaviors within aggregates. It is validated against two gold-standard bulk assays performed on spherical aggregates: micropipette aspiration and Hertzian plate compression. Under aspiration, the model demonstrates a transition from elastic deformation to viscous creep governed by localized packing and emergent jamming at the aspirated neck, accompanied by increased mechanically adaptive remodeling. Under compression, core rheology determines deformation mode: liquid-like aggregates exhibit enhanced swirling, consistent with experimental observations, whereas solid-like aggregates exhibit affine, Poisson-like deformation. These results bridge cell-scale dynamics and quantifiable tissue rheology including elastic modulus and vicosity, providing a framework to interpret emerging 3D measurements of multicellular mechanics.

## INTRODUCTION

Multicellular tissues behave as complex mechanical materials whose macroscopic rheology emerges from interactions between individual cells. Tissues transition between elastic deformation, viscous flow, collective rearrangement, and jamming depending on mechanical loading and cellular activity [1–5], during morphogenesis [6, 7], wound healing [8], and tumor progression [9]. Understanding how these emergent behaviors arise from cell-level interactions remains a central challenge in biophysics and developmental mechanics.

At the single-cell level, mechanical behavior is governed by cytoskeletal networks and cell–cell junctions that generate and transmit forces across tissues [8–10], capable of dissipating energy, transmitting tension, and remodeling under deformation [11, 12]. Crucially, cell–cell junctions are highly dynamic structures that continuously remodel through cytoskeletal contractility, adhesion turnover, and mechanotransduction, enabling local interactions to reorganize in response to mechanical cues [8, 10].

At the tissue scale, multicellular aggregates exhibit rate-dependent viscoelastic behavior [1], transitioning from elastic deformation at short times to viscous flow under sustained loading [2]. These bulk properties are commonly quantified using experimental assays such as micropipette aspiration [2, 13, 14], compression testing [1, 15, 16], and tensile testing [17, 18]. Recent studies further suggest that tissue mechanical states are governed by cell-level structural transitions, including rigidity percolation and connectivity-driven phase behavior [3, 5, 19]. Our prior work, Tomizawa et al. [20], extends these observations to single-cell resolution within intact tissues, revealing heterogeneous solid-like and liquid-like mechanical responses at the cellular level [20]. Together, these findings highlight the need for frameworks that derive tissue rheology directly from cell–cell interactions.

Several theoretical frameworks have been developed to model these behaviors. Continuum finite-element approaches capture macroscopic stress–strain responses and enable quantitative interpretation of bulk mechanical assays [13, 14, 20]. However, they prescribe constitutive laws defined at the tissue scale. As a result, complex collective behaviors, such as swirling flows, cell topology rearrangements, and turnover observed in active tissues [21] must be introduced via additional modeling layers [22, 23], rather than emerging directly from underlying cell–cell interactions.

In contrast, discrete cell-based models explicitly represent individual cells and their interactions. Vertex models describe tissues as networks of polyhedral cells governed by cortical tension, adhesion, and volume elasticity [24, 25], while Cellular Potts models employ lattice-based Monte Carlo dynamics to capture processes such as cell sorting, morphogenesis, and angiogenesis [26]. These approaches demonstrate how geometric constraints and connectivity give rise to collective phenomena including jamming and rigidity transitions [25, 27, 28]. However, their standard formulations emphasize energy minimization and local mechanics rather than bulk rheology. Although they can exhibit viscoelastic behavior, incorporating mechanosensitive junction remodeling [10, 11] typically requires extensions beyond their simplest implementations [29–31].

This highlights a gap in current modeling approaches in relating dynamic cell-level interactions to macroscopic rheology and the mechanisms underlying complex viscoelastic behavior. Continuum frameworks prescribe tissue-scale rheology without resolving the underlying cell–cell interactions, whereas discrete models capture cell-level organization and rearrangements but do not directly link these interactions to emergent tissue-scale mechanical response. This motivates a framework in which tissue rheology arises directly from single-cell interactions, consistent with experimental evidence that collective mechanical behavior emerges from junction-level dynamics and remodeling [10, 11].

Here, we introduce a three-dimensional network model in which each cell–cell connection acts as a viscoelastic element that evolves with its mechanical history. Supporting this perspective on the molecular scale, the Gaussian Network Model demonstrates how collective protein dynamics can be inferred directly from residue–residue interaction topology, with amino acids represented as nodes in an elastic network whose connectivity determines large-scale fluctuations and correlated motions [32]. Applying an analogous network-based approach for multicellular tissues, with cells represented as nodes in a viscoelastic network whose interaction topology gives rise to tissue-scale mechanical behavior, remains largely unexplored.

We will show that macroscopic rheology emerges from these adaptive, history-dependent interactions, rather than from prescribed bulk parameters. A key innovation is a remodeling mechanism that governs both the mechanical strength and rest length of connections in response to local deformation. Using in-silico realizations of two standard experimental assays on spherical aggregates, micropipette aspiration [2] and Hertzian compression [15], we aim to bridge single-cell mechanics and bulk tissue rheology, providing insight into the local behaviors that govern macroscopic response.

## MODEL OVERVIEW

### Network Construction

A multicellular aggregate is represented as a network of *N* cells modeled as nodes in ℝ^3^. Each node *i* ∈ {1, …, *N}* has position **x**_*i*_(*t*) corresponding to the center of a single cell. The initial configuration {**x**_*i*_(0)} is generated within a spherical domain of radius *R*, representing a multicellular aggregate (Fig. 1).

**FIG. 1.**
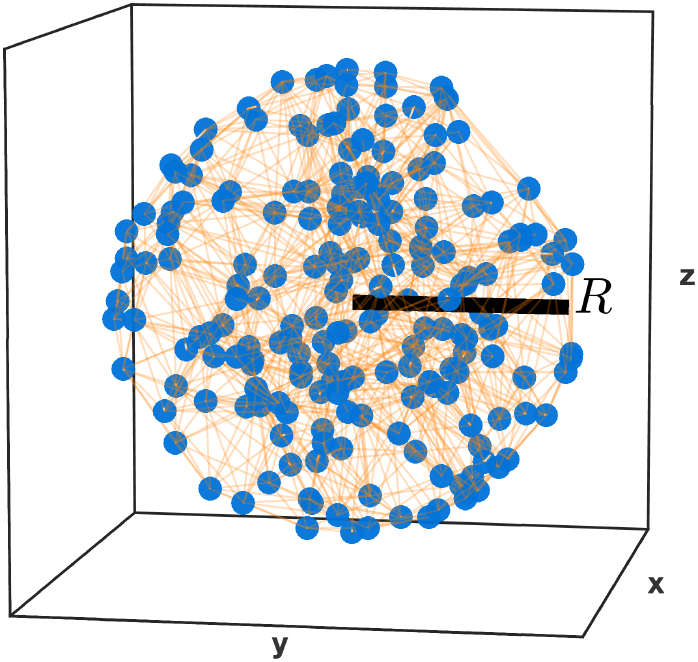
Spherical aggregate. 3D Aggregate with *N* = 200 cells and radius *R*. Cell centers (blue) are connected to their neighbors via viscoelastic elements (orange). Simulated aggregates typically contained *N* = 500 cells.

Mechanical interactions are encoded by an undirected graph *G* = (*V*, ℰ), where *V* denotes nodes and ℰ connections. Connectivity is defined using a three-dimensional Delaunay triangulation of the initial node positions, creating an irregular but space-filling structure similar to how cells are packed together in multicellular aggregates.

For each connection *e* = (*i, j*) ∈ ℰ, the instantaneous separation between cells is

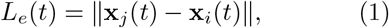

with unit direction

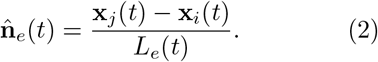

The initial rest length of each connection is defined as

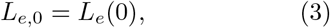

which serves as the reference length prior to mechanical loading. As described in following sections, these rest lengths may evolve dynamically through mechanically adaptive remodeling.

Mechanical coupling between neighboring cells is modeled at the level of individual network connections. Each connection *e* ∈ ℰ is treated as a 1D Burgers viscoelastic element acting along the line connecting two cell centers (Fig. 2). This formulation assigns viscoelasticity directly to cell–cell interactions themselves, allowing heterogeneous and history-dependent mechanical responses to emerge naturally across the network.

**FIG. 2.**
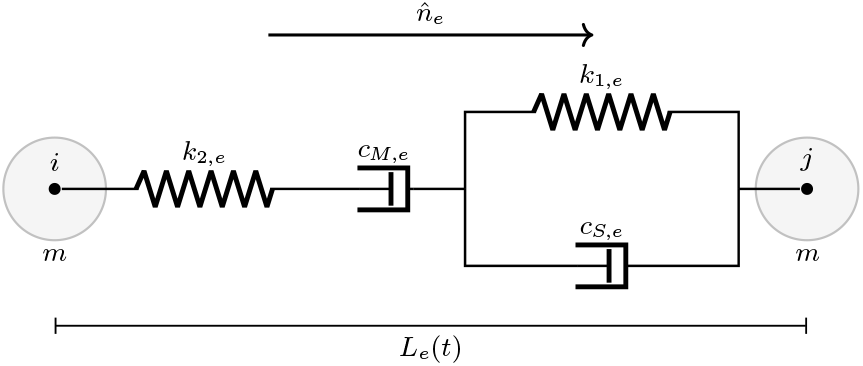
Cell-cell viscoelastic connection schematic. Burgers element connecting neighboring cells *i* and *j* of mass *m* and length *L*_*e*_(*t*) with reference unit direction 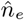. The connection consists of a Maxwell element (*k*_2,*e*_ and *c*_*M,e*_) in series with a Kelvin–Voigt element (*k*_1,*e*_ and *c*_*S,e*_).

For a connection *e* = (*i, j*), let *L*_*e*_(*t*) denote its instantaneous length and *L*_0,*e*_(*t*) its adaptive rest length. The extension and corresponding dimensionless strain are defined as

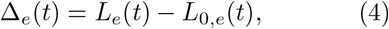

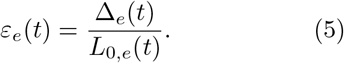

The rate of extension along the connection direction is given by

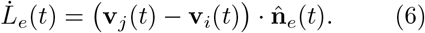

The four viscoelastic parameters *k*_1,*e*_, *k*_2,*e*_, *c*_*S,e*_, and *c*_*M,e*_ are dynamic and may change over time as a result of mechanical adaptation. The internal dynamics are described by the connection tension *σ*_*e*_(*t*) and a Kelvin–Voigt extension *z*_*e*_(*t*),

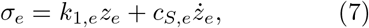

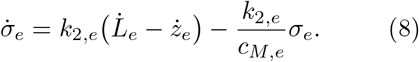

The resulting connection force acts along the line of centers,

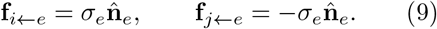

Node forces are obtained by summing contributions from all incident connections.

### Mechanically Adaptive Remodeling

Cell–cell junctions are inherently dynamic, continuously remodeling through cytoskeletal contractility, adhesion turnover, and mechanotransduction [8, 10]. To account for this, we introduce a remodeling framework in which intercellular interactions evolve in response to mechanical cues.

The mechanical history of each cell-cell connection is encoded through memory variables for sustained proximity of neighboring cells, tension, and compression,

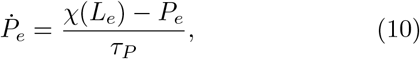

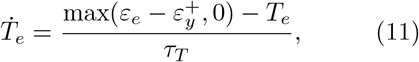

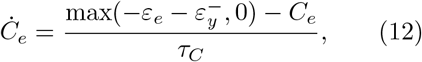

where *χ*(*L*_*e*_) is a proximity kernel and *τ*_*P*_, *τ*_*T*_, *τ*_*C*_ are characteristic memory times. The yield strains 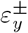 ensure that only sustained suprathreshold deformation contributes to remodeling.

These variables act as low-pass filters of the local mechanical state, allowing each connection to accumulate a memory of persistent proximity, tension, or compression over time rather than responding instantaneously. In this sense, the remodeling dynamics “lag behind” the instantaneous deformation, approaching the current mechanical state over characteristic timescales. As a result, only sustained mechanical signals, rather than transient fluctuations, drive changes in intercellular properties.

Each connection carries a weight coefficient *w*_*e*_(*t*) that scales its viscoelastic parameters *k*_1,*e*_, *k*_2,*e*_, *c*_*S,e*_, and *c*_*M,e*_. The target state is determined by a remodeling signal

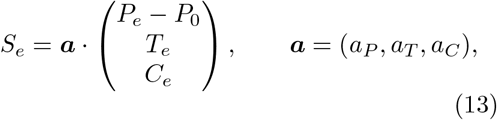

where *P*_0_ is a reference proximity level corresponding to neutral adhesion remodeling. The coefficients a = (*a*_*P*_, *a*_*T*_, *a*_*C*_) weight the relative contributions of sustained proximity, tensile loading, and compressive loading to the remodeling response. The weight relaxes toward

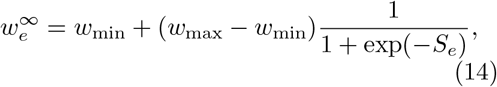

allowing connections to strengthen or weaken depending on sustained mechanical cues. The smooth sigmoidal form avoids abrupt transitions while capturing the intrinsically dynamic nature of cell–cell junctions. In this formulation, weakening of connections (*w*_*e*_ → 0) reflects junctional turnover and dissociation, enabling neighbor exchange during processes such as collective cell migration and invasion [33], while strengthening (*w*_*e*_ *>* 1) captures mechanosensitive reinforcement of connections under load. Together, these dynamics allow the interaction network to continuously reorganize, consistent with experimentally observed junctional plasticity and force-dependent remodeling [6, 8, 11].

Rest lengths evolve irreversibly when strain exceeds yield thresholds, allowing relaxation of accumulated strain. The plastic evolution follows

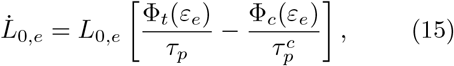

where Φ_*t*_ and Φ_*c*_ are smooth functions that activate only beyond tensile or compressive yield thresholds and increase with excess strain. Lower bounds on *L*_0,*e*_ prevent geometric collapse. This formulation adopts a Perzyna-type thresholded viscoplastic structure, in which irreversible deformation is activated beyond yield and increases smoothly with excess deformation [34, 35], consistent with experimentally observed yielding in cells and tissues [36].

The characteristic remodeling timescales (*τ*_*P*_, *τ*_*T*_, *τ*_*C*_, *τ*_*p*_) are chosen to be consistent with experimentally observed timescales for actomyosin turnover and cell–cell junction remodeling, which occur on the order of minutes to hours and govern stress dissipation and viscoelastic behavior in tissues [8, 11, 37]. Together, these mechanisms allow each connection to adapt its strength and geometry in response to sustained deformation, enabling long-duration mechanical adaptation.

## MICROPIPETTE ASPIRATION

### Aspiration Dynamics

To evaluate bulk rheology, we simulated micropipette aspiration of a spherical aggregate of *N* = 500 cells with initial radius *R*_0_ = 125 *µ*m. Aspiration was imposed through a cylindrical pipette of radius *R*_*p*_ = 35 *µ*m under a pressure difference Δ*P* = 1.2 kPa. Following Guevorkian *et al*. [2], surface tension introduces a critical pressure Δ*P*_*c*_ = 400 Pa, giving an effective driving pressure Δ*P* − Δ*P*_*c*_ = 800 Pa. The resulting aspiration force is

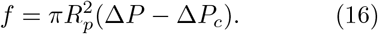

The aspiration length *L*(*t*) was computed from leading cells in within the pipette cylinder relative to the mouth plane, after tongue formation. To interpret the emergent aspiration dynamics, we analyze the simulated aspiration length *L*(*t*) using the modified Maxwell framework introduced by Guevorkian et al. [2],

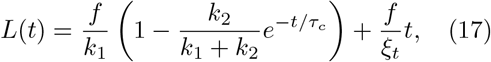

where *k*_1_ and *k*_2_ are elastic constants, *τ*_*c*_ is the transient relaxation time, and *ξ*_*t*_ is the effective viscous drag. Importantly, the modified Maxwell model is not imposed during simulation, but is instead used only to extract equivalent continuum parameters from the aspiration response.

For fitting, we recast this expression in the observable form

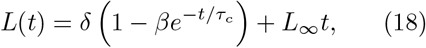

with

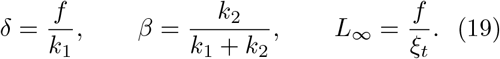

Fitting yields *δ* = 2.654 × 10^−5^ m, *L*_∞_ = 8.571 × 10^−9^ m*/*s, *β* = 0.5, and *τ*_*c*_ = 993 s, implying *k*_1_ ≈ *k*_2_. Using

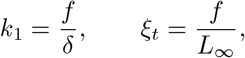

and 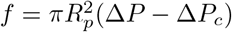 gives

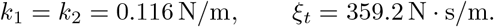

Following [2], these coefficients relate to elastic modulus *E* and viscosity *η* via

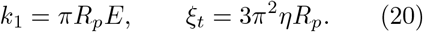

Using *R*_*p*_ = 35 *µ*m yields

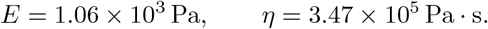

These values are consistent with reported ranges for multicellular spherical aggregates [2, 38], demonstrating the emergence of bulk tissue rheology from the underlying cell-level mechanics.

The elastic–viscous crossover is governed by the Maxwell time

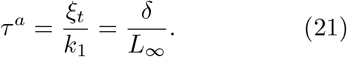

Using the fitted parameters gives *τ* ^*a*^ = 3.10 × 10^3^ s ≈ 51.6 min. In reference to the onset of aspiration after tongue formation, the effective crossover time is 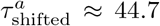 min, consistent with experimental relaxation times [2]. Notably *τ* ^*a*^ ≈ 3 *τ*_*c*_, indicating a separation between the fast elastic relaxation (*τ*_*c*_) and the slower viscous creep. The dashed vertical line in Fig. 3(d,e) marks this crossover.

**FIG. 3.**
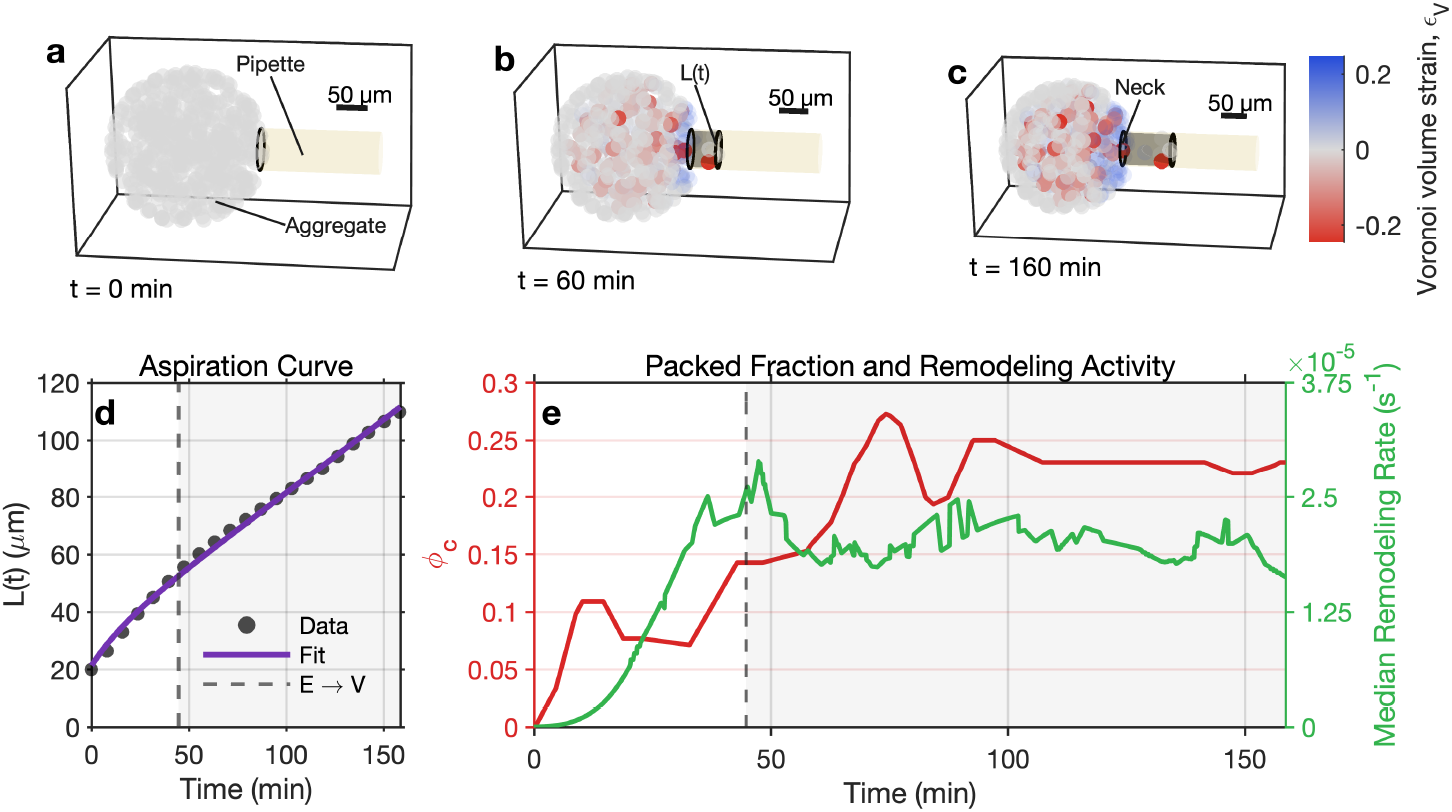
Aspiration-induced deformation reveals emergent jamming and mechanically adaptive remodeling in a 3D viscoelastic cell network. (a–c) 3D snapshots of a spherical aggregate undergoing micropipette aspiration [2] at *t* = 0, 60, and 160 min, with pipette shading indicating aspiration length. Cells are colored by Voronoi volume strain (*ϵ*_*V*_), showing local compaction (red) and dilation (blue). Deformation localizes at the pipette neck, where compression and structural reorganization concentrate. **(d)** Aspiration length *L*(*t*) (*µ*m) showing the transition from elastic deformation to viscous creep. Black circles indicate simulation data; the solid line is the viscoelastic model fit (*R*^2^ = 0.998). The dashed vertical line marks the elastic-to-viscous (E → V) transition. **(e)** Neck-region packed fraction (*ϕ*_*c*_, red, left axis), used as a proxy for jamming, and median remodeling rate (green, right axis) over time. The dashed vertical line again marks the E → V transition, where packing and remodeling increase, indicating jamming and load-dependent adaptation of intercellular connections.

### Neck-Localized Packing, Emergent Jamming, and Remodeling

We note that mechanical loading during aspiration is spatially heterogeneous: the pipette entrance concentrates stress transmission be-(21) tween the aspirated tongue and the aggregate, creating a confined region in which compression and load-bearing interactions are amplified. We thus quantify structural evolution within a cylindrical “neck” region centered on the pipette mouth (Fig. 3c).

Local deformation is characterized using the Voronoi volume strain

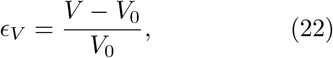

where *V* is the instantaneous Voronoi cell volume and *V*_0_ its reference value. Because Voronoi volumes depend only on instantaneous cell-center geometry, *ϵ*_*V*_ provides a single-cell-resolved measure of local packing compatible with the Delaunay interaction network governing force transmission. Coloring cells by *ϵ*_*V*_ (Fig. 3a–c) reveals progressive localization of compression at the aspirated neck. As aspiration proceeds, volumetric strain becomes increasingly negative in this region while the aggregate shell remains comparatively less compressed. To quantify this effect we define a neck packed fraction

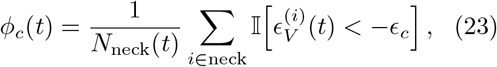

where I[·] is the indicator function and *ϵ*_*c*_ is a compression threshold. As shown in Fig. 3e, *ϕ*_*c*_ increases markedly during aspiration, indicating growing local crowding consistent with jamming at the pipette entrance. Notably, the rise in *ϕ*_*c*_ occurs near the independently determined crossover time *τ* ^*a*^, indicating that the transition from elastic deformation to viscous creep coincides with the formation of a jammed, mechanically constrained neck that regulates subsequent flow into the pipette.

Because viscoelastic interactions are coupled to adaptive remodeling, we also quantify neck-localized remodeling activity through the median connection-weight evolution rate,

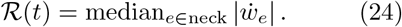

Remodeling activity increases in tandem with *ϕ*_*c*_ (Fig. 3e), indicating that the region undergoing compression also exhibits enhanced adaptation of cell–cell connections.

We further observe a temporally ordered shift in remodeling direction in the neck region. During early elastic deformation, we observe that most cell–cell connections tend to weaken 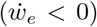, consistent with initial yielding of the network under applied stress. During viscous creep, cell–cell connections strengthen 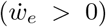, indicating the formation of a new mechanically stable configuration. The E → V transition therefore marks the point at which this shift occurs.

In addition, because the aggregate is represented as a finite collection of cells, the model captures a gradual shrinkage of the reservoir below the pipette as cells are drawn inward. Although modest over the simulated time window, this effect reflects a finite geometric constraint absent in continuum models that assume an effectively infinite reservoir [2]. Over sufficiently long aspiration times, conservation of mass implies that the entire aggregate would ultimately be drawn into the pipette, with the reservoir volume vanishing.

## HERTZIAN PLATE COMPRESSION

To probe whether the model reproduces experimentally observed short-term deformation modes, we performed quasi-static Hertzian plate compression simulations and analyzed both the macroscopic force–indentation response and the resulting single-cell displacement fields (Fig. 4). Because Hertzian compression is applied over a short timescale (seconds), mechanically adaptive remodeling is effectively inactive during this assay. The characteristic remodeling timescales (*τ*_*P*_, *τ*_*T*_, *τ*_*C*_, *τ*_*p*_) are orders of magnitude longer, such that connection weights *w*_*e*_(*t*) and rest lengths *L*_0,*e*_(*t*) remain approximately constant throughout compression. The response thus reflects the instantaneous viscoelastic properties of the intercellular connections.

**FIG. 4.**
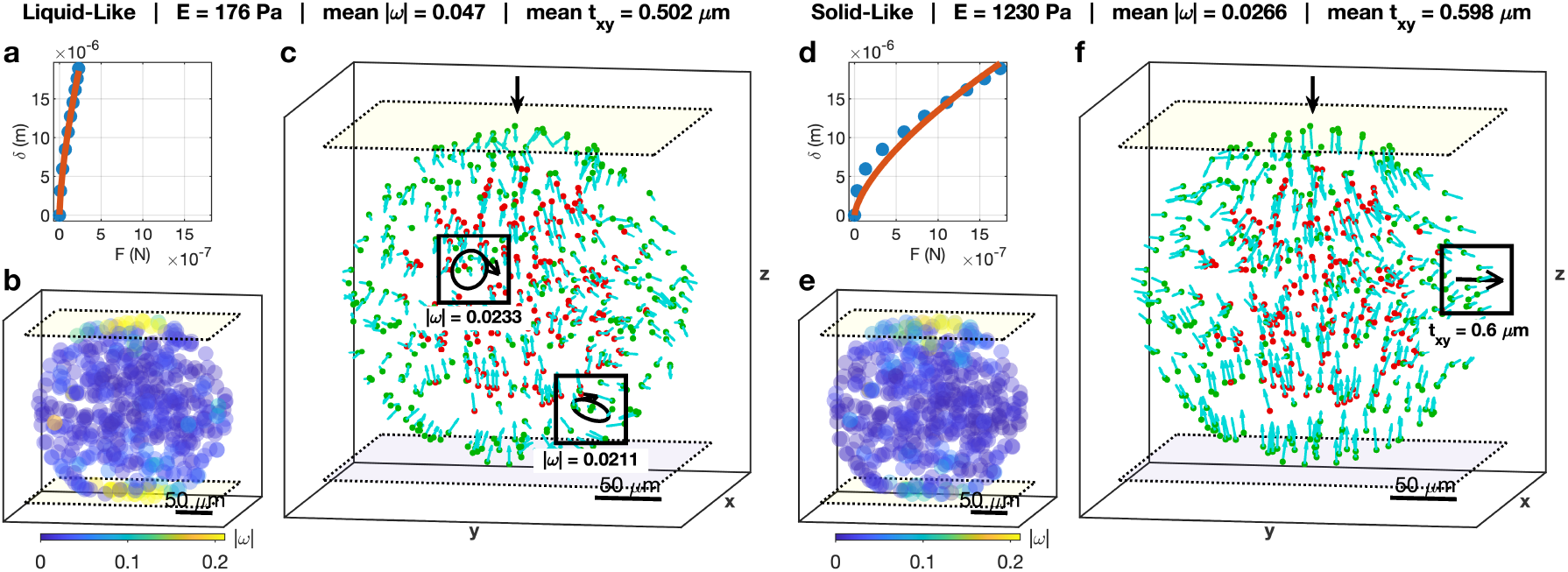
Emergent rotational dynamics versus Poisson-like deformation in *in silico* spherical aggregates under Hertzian compression. **(a,d)** Force–indentation curves from compression of two simulated spherical aggregates. Blue dots show simulation data and red lines show Hertzian fits. The liquid-like aggregate has an effective modulus *E* ≈ 176 Pa (*R*^2^ = 0.998), while the solid-like aggregate yields *E* ≈ 1230 Pa (*R*^2^ = 0.962), indicating distinct bulk moduli arising from cell–cell rheology. **(b**,**e)** Color maps of swirling magnitude |*ω*| during compression. The liquid-like aggregate shows elevated rotational activity compared to the solid-like aggregate (*p* ≪ 0.001), consistent with experimental observations [20]. **(c**,**f)** Cellular displacement fields during compression (cyan arrows). Cells are colored by shell membership (red = inner core, green = outer shell). **(c)** In the liquid-like aggregate, single cell displacements organize into rotational swirling motions (boxed regions). **(f)** In the solid-like aggregate, displacements are aligned with the compression direction, showing greater Poisson-like lateral expansion (*t*_*xy*_, boxed region) compared to the liquid-like aggregate (*p* ≪ 0.001).

This assay mirrors our previous Hertzian compression analyses [15] and recent light-sheet micromechanical compression experiments on composite organoids composed of trophoblast and stromal fibroblast cells [20], which organize into a core–shell architecture respectively with distinct mechanical responses. In these experiments, fibroblast-rich regions exhibited either solid-like or liquid-like behavior depending on their state. Solid-like regions displayed well-aligned axial compression with Poisson-like lateral expansion, whereas liquid-like regions underwent disordered cellular rearrangements with swirling and rotational motion. Here, we simulate two spherical aggregates with identical geometry, topology, and initial configuration, such that the only difference is the rheological state of the core (liquid-like versus solid-like), achieved by altering the spring constants and damping coefficients of the viscoelastic arms.

### Bulk Moduli via Hertzian Fitting

The macroscopic response is characterized by the plate reaction force *F* as a function of indentation depth *δ*. Under quasi-static loading, the force–indentation relation follows the Hertz contact law for a spherical body,

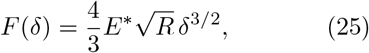

where *R* is an effective radius of curvature and

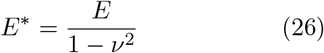

is the effective modulus.

To estimate *E*, we linearize Eq. (25) as

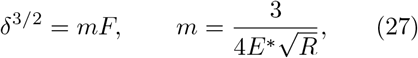

and determine *m* by least-squares regression. The effective Young’s modulus then follows as

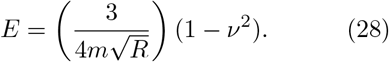

Applying Eq. (28) with *ν* = 0.49 gives distinct bulk moduli for the two simulated aggregates: *E* ≈ 176 Pa for the liquid-like aggregate and *E* ≈ 1230 Pa for the solid-like aggregate (Fig. 4a,d).

### Single-cell Swirling Metric

To distinguish aligned (solid-like) deformation from rotational (liquid-like) motion observed experimentally under compression [20], we compute a node-resolved swirling magnitude |ω_*i*_| from the displacement field using a local curl-like estimator.

Let **x**_*i*_ denote the position of node *i* and **u**_*i*_ its displacement relative to the reference configuration. For each node we define a fixed set of *k* nearest neighbors *N*_*i*_ based on the reference positions. For *j* ∈ *N*_*i*_, the relative position and displacement are

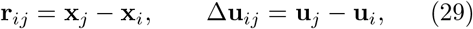

with radial weights that emphasize nearby neighbors,

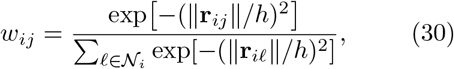

where *h* is a characteristic nearest-neighbor length scale. The local curl-like vector is then

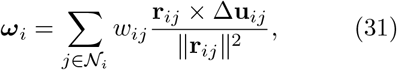

with magnitude

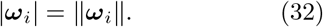

This local kinematic estimator isolates rotational motion while suppressing uniform translation. Color maps of |ω_*i*_| are shown in Fig. 4b,e.

### Rotational Dynamics versus Poisson-like Deformation

Fig. 4 summarizes the key qualitative and quantitative outcomes. Both simulated spherical aggregates follow Hertz-like force– indentation curves (Fig. 4a,d), but their deformation fields differ markedly. To quantify these differences, we analyzed single-cell rotational magnitude |ω| and lateral displacement *t*_*xy*_. Statistical comparisons reveal systematic shifts in central tendency between regimes: liquid-like aggregates exhibit higher rotational activity (Fig. 4b, c) and reduced lateral translation, whereas solid-like aggregates exhibit the opposite trend (Fig. 4e,f), with *p* ≪ 0.001 for both metrics.

These trends mirror our experimental observations under quasi-static micro-compression, where solid-like regions exhibited well-aligned displacement vectors while liquid-like regions displayed eddy-like swirling and rotational motion [20]. In experiments, liquid-like behavior correlated with elevated von Mises strain [20]; here, the |ω| maps provide a complementary single-cell-resolved measure that isolates rotational components of the displacement field. Applying the same swirl formulation to experimental single-cell tracking data from our previous work, Tomizawa et al. [20], reveals the same statistical trend (*p* ≪ 0.001) and comparable magnitudes of rotational activity, with liquid-like aggregates exhibiting enhanced swirling relative to solid-like counterparts.

## DISCUSSION AND CONCLUSION

We have shown a three-dimensional mechanically adaptive viscoelastic cell-network model in which tissue-scale rheology emerges directly from cell–cell interactions and their history-dependent remodeling. Our model uniquely highlights single-cell behaviors such as rotational swirling and localized jamming that govern the bulk mechanical response, while achieving agreement with experimentally reported ranges for elastic modulus [15] and viscosity [2, 38] for spherical aggregates.

Under Hertzian compression, the model reveals distinct deformation modes: liquid-like aggregates exhibit rotational, swirling motion, whereas solid-like aggregates deform in a more affine, Poisson-like manner, consistent with our single-cell-resolved experiments [20] and observations of collective dynamics in active tissues [21]. Our method therefore enables prediction and quantification of both mechanical state and progression through phase-like transitions, including epithelial–mesenchymal (EMT) [39, 40], mesenchymal–epithelial (MET) [41], and decidualization [20], from experimental observations.

In micropipette aspiration, the transition from elastic deformation to viscous creep coincides with the emergence of a localized, load-bearing neck characterized by increased jamming and elevated remodeling activity. This result suggests that the elastic–to-viscous crossover may be characterized not only as a bulk material property but also as a microscale structural transition [19]. We therefore predict that experimental perturbations altering adhesion turnover [42], actomyosin remodeling [43], or effective cell–cell connectivity [4, 19, 25, 28] can shift the onset of local crowding and the elastic-to-viscous crossover, making these structural signatures directly testable in aspiration experiments.

Furthermore, our remodeling mechanism naturally captures history-dependent behavior observed in biological tissues [11, 37], consistent with emerging models of adaptive viscoelasticity and mechanical memory [29, 31]. Since each cell–cell connection carries its own dynamically evolving mechanical properties and memory of past interactions, repeated or prolonged loading alters subsequent mechanical response both locally and at the tissue scale. As such, we note that in our model, cells effectively “communicate” with one another through their viscoelastic connections. Our approach therefore provides a natural foundation for incorporating network-based biochemical communication [44], allowing cell-cell signaling pathways regulating adhesion and contractility to directly modulate tissue-scale mechanics [8, 10].

In summary, our model bridges cell-scale dynamics and tissue rheology. It presents a framework for interpreting emerging 3D, single-cell resolved measurements of multicellular mechanics and provides insight into the single-cell behaviors that influence bulk tissue response.

## Supporting information

Compression Motion Solid Like

Compression Motion Liquid Like

